# Astrocytes mediate a positive feedback loop for oxytocin

**DOI:** 10.64898/2026.02.02.699227

**Authors:** Maria Clara Selles, Melissa L. Cooper, Francesco Limone, Araf Ahmed, Shane A. Liddelow, Robert C. Froemke, Moses V. Chao

## Abstract

Social interactions are critical for well-being and survival. Oxytocin neurons in the paraventricular nucleus of the hypothalamus help regulate social behaviors in many species, and respond to social stimuli to promote pro-social interactions. Here, we show that chronic social isolation reduced production of oxytocin peptide, and led to a delay in the onset of huddling behavior upon resocialization in male mice. Exogenous oxytocin treatment prevented both the behavioral and molecular effects of social deprivation. Using conditional knockouts, we found that oxytocin-induced oxytocin expression was mediated by local hypothalamic astrocytes. Oxytocin signaling in astrocytes upregulated the expression of a retinoic acid-synthesizing enzyme Aldh1a1, and retinoic acid increased oxytocin expression. These findings reveal a mechanism in which astrocytes can sense and control neuropeptide levels to influence social behaviors.

## Main text

Social behaviors are necessary for survival of vertebrate species (*1*). Social interactions depend on two main factors: 1) acute sensory inputs, including the environment and social cues from other animals; and 2) internal states related to the animal performing the behavior (*2*). In the context of an epidemic of loneliness and social isolation (*3*), understanding the effects of social deprivation on internal states and its behavioral consequences becomes an urgent matter to develop more effective prevention strategies and treatment approaches.

Neuropeptides, such as oxytocin, have been shown to play central roles by sensing social context and modulating social behaviors (*4–9*). Oxytocin activates oxytocin receptors (OXTRs), which are expressed in neuronal and glial cells (*10*). Astrocytes, a type of glial cell, play a main role in neuronal homeostasis (*11*). They have been shown to express OXTRs and to induce behavioral changes in response to oxytocin (*12*). Moreover, recent work shows that astrocytes present context-dependent responses to oxytocin and other neurotransmitters, highlighting the relevance of internal states not only for neuronal but also for astrocyte function (*13, 14*).

Here we examined the effects of long-term social isolation on the oxytocin system and its consequences for affiliative social behaviors in mice. We aimed to understand how neurons and astrocytes communicate in the context of social isolation to modulate future social encounters. We describe the molecular mediators of a novel hypothalamic positive feedback loop in which oxytocin upregulates its own expression in an astrocyte-dependent manner, and its relevance for social behaviors.

## Results

### Chronic social isolation reduces oxytocin production and delays huddling in male mice

To study the behavioral and molecular effects of chronic social deprivation, we isolated adult male mice for four weeks. We tested the effect of long-term social isolation on social behaviors in a reunion paradigm taking advantage of long-term 24/7 continuous behavioral monitoring (*15*), and on oxytocin mRNA and peptide levels using qPCR and iDISCO immunolabeling (**Fig. 1A**). Isolated mice were compared to group-housed control age-matched males. We found that isolated mice exhibited a delay in huddling onset when paired with another animal compared to the huddling onset time in animals that were group housed (**Fig. 1B**, p=0.0007). In separate cohorts of mice, we found that social isolation downregulated oxytocin production at the mRNA (**Fig. 1C**, p=0.0498) and peptide levels (**Fig. 1D**, p=0.0096). We used Imaris to reconstruct oxytocin neurons in the PVN, and observed a reduction of oxytocin-positive PVN volume in mice that were socially isolated (**Fig. 1E**, p=0.0265).

**Fig. 1.**
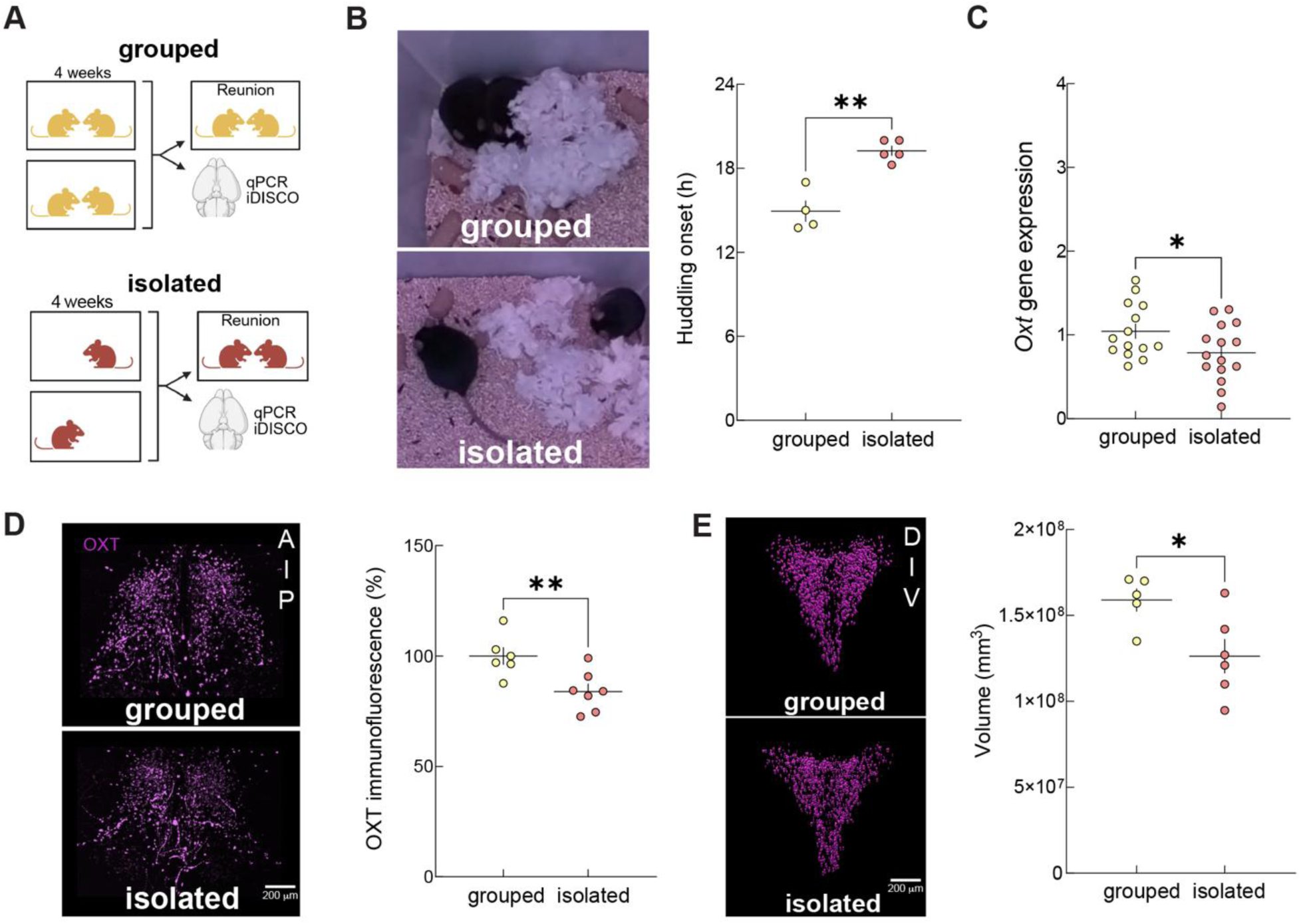
Behavioral and molecular effects of chronic social isolation. (**A**) Schematic representation of the 4 weeks long social isolation protocol, followed by behavioral analysis upon reunion with a previous littermate, or molecular characterization of oxytocin levels in the hypothalamus. (**B**) Representative image and quantification of the latency to start huddling upon reunion. (**C**) qPCR quantification of hypothalamic oxytocin mRNA levels. (**D**) Maximum intensity projection of PVN z-stacks in iDISCO-immunolabeled and cleared brains and quantification of oxytocin peptide levels by immunofluorescence. (**E**) Imaris reconstruction of oxytocin positive cells in the PVN and volume quantification of the region. Each dot represents a mouse or a mouse pair in reunion experiments. All error bars represent SEM. Unpaired two-tailed Student’s t-test. *, p<0.05; **, p<0.01.

### Oxytocin upregulates its own expression in socially isolated male mice

We next asked if the behavioral phenotype observed after social deprivation (slower emergence of huddling) could be mediated by a decrease in oxytocin levels. We used two methods to elevate oxytocin levels: intranasal delivery of exogenous oxytocin (**Fig. 2A,B**) and chemogenetics to increase endogenous release from the PVN (**Fig. 2C,D**).

**Fig. 2.**
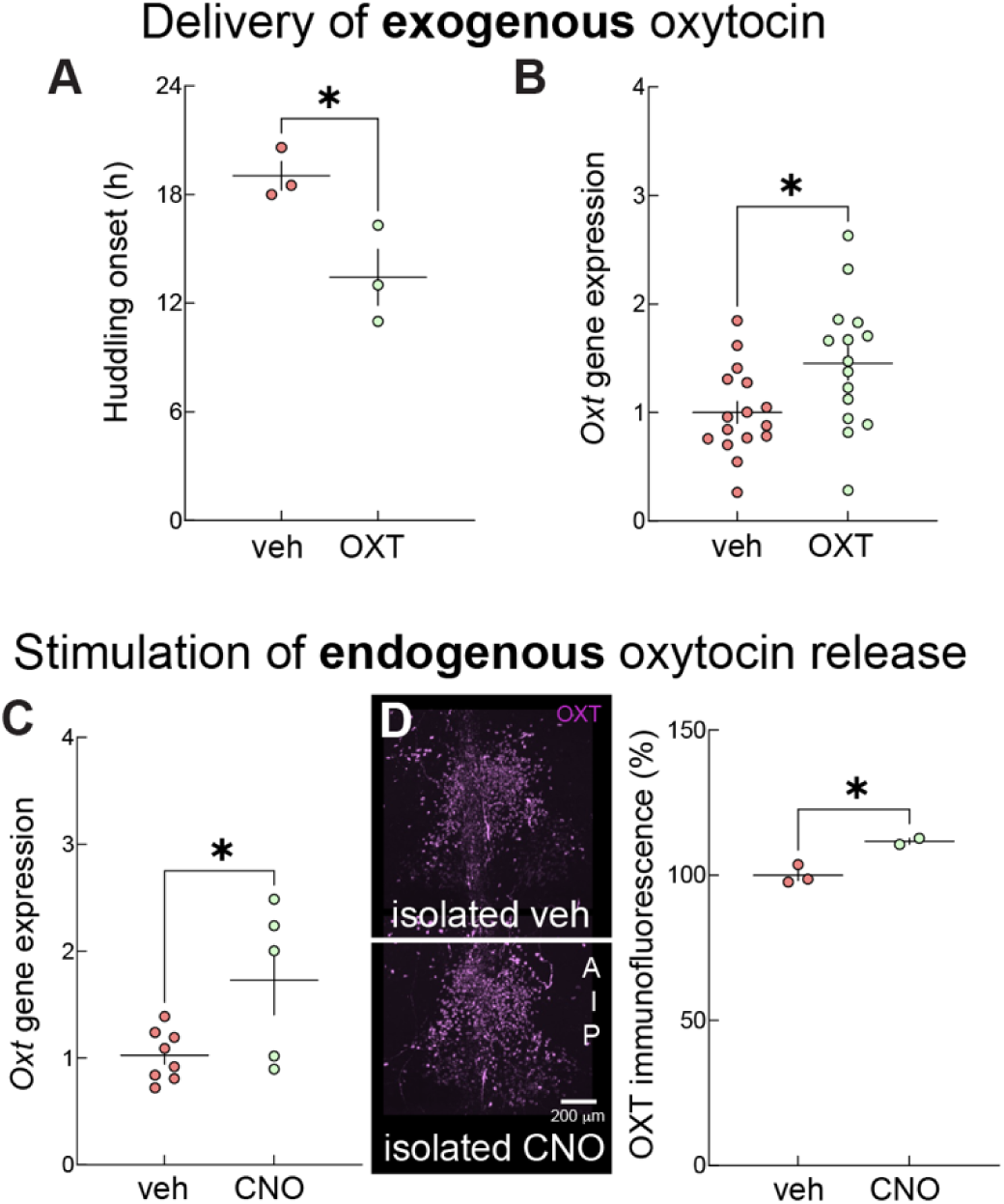
A novel positive feedback loop for oxytocin. (**A**) Quantification of the latency to start huddling upon reunion and (**B**) oxytocin mRNA levels in mice that received intranasal oxytocin (or vehicle as control) three times a week during the 4 weeks of isolation. (**C**) Quantification of oxytocin mRNA and (**D**) peptide levels in Oxy-Cre mice bilaterally injected with an AAV-hSyn-DIO-hM3D(Gq) virus in the PVN and implanted with a subcutaneous osmotic pump to release CNO (or vehicle as control) during the 4 weeks of isolation. Each dot represents a mouse or a mouse pair in reunion experiments. All error bars represent SEM. Unpaired two-tailed Student’s t-test, *, p<0.05.

For the exogenous approach, we delivered intranasal oxytocin three times per week during the 4 week-long isolation period and found that it prevented the delay in huddling onset induced by social deprivation (**Fig. 2A**, p=0.0322). We tested whether exogenous oxytocin delivery could impact the endogenous system, and found that it upregulated endogenous oxytocin production (**Fig. 2B**, p=0.0198).

To test whether stimulation of the endogenous system could have a similar effect on oxytocin production, we used Oxytocin-Ires Cre (Oxy-Cre) mice injected with an AAV-hSyn-DIO-hM3D(Gq) virus. We injected the virus bilaterally in the PVN at the age of 1.5 months (two weeks before social isolation) to express excitatory Designer Receptor Exclusively Activated by Designer Drugs (DREADDs) in oxytocin neurons. At the age of 2 months, mice were implanted subcutaneously with Alzet osmotic pumps containing either clozapine N-oxide (CNO) -the DREADD ligand- or vehicle, and socially isolated for a month-long period. We found that chemogenetic stimulation of oxytocin neurons led to upregulation of oxytocin expression at the mRNA (**Fig. 2C**, p=0.0253) and peptide levels (**Fig. 2D**, p=0.0189).

### Astrocytes mediate a positive feedback loop for oxytocin

To investigate the cellular mechanisms by which oxytocin can increase its own expression, we focused on astrocytes as these glial cells are key mediators of neuronal plasticity (*16*), express OXTRs, and respond to oxytocin (*12*). We crossed transgenic Aldh1l1-Cre/ERT2 mice with OXTR^fl/fl^ mice to knock-out OXTRs from astrocytes (**Fig. 3A**). We found that mice lacking OXTRs in astrocytes fail to increase endogenous oxytocin production in response to exogenous treatment (**Fig. 3B**, p=0.0808).

**Fig. 3.**
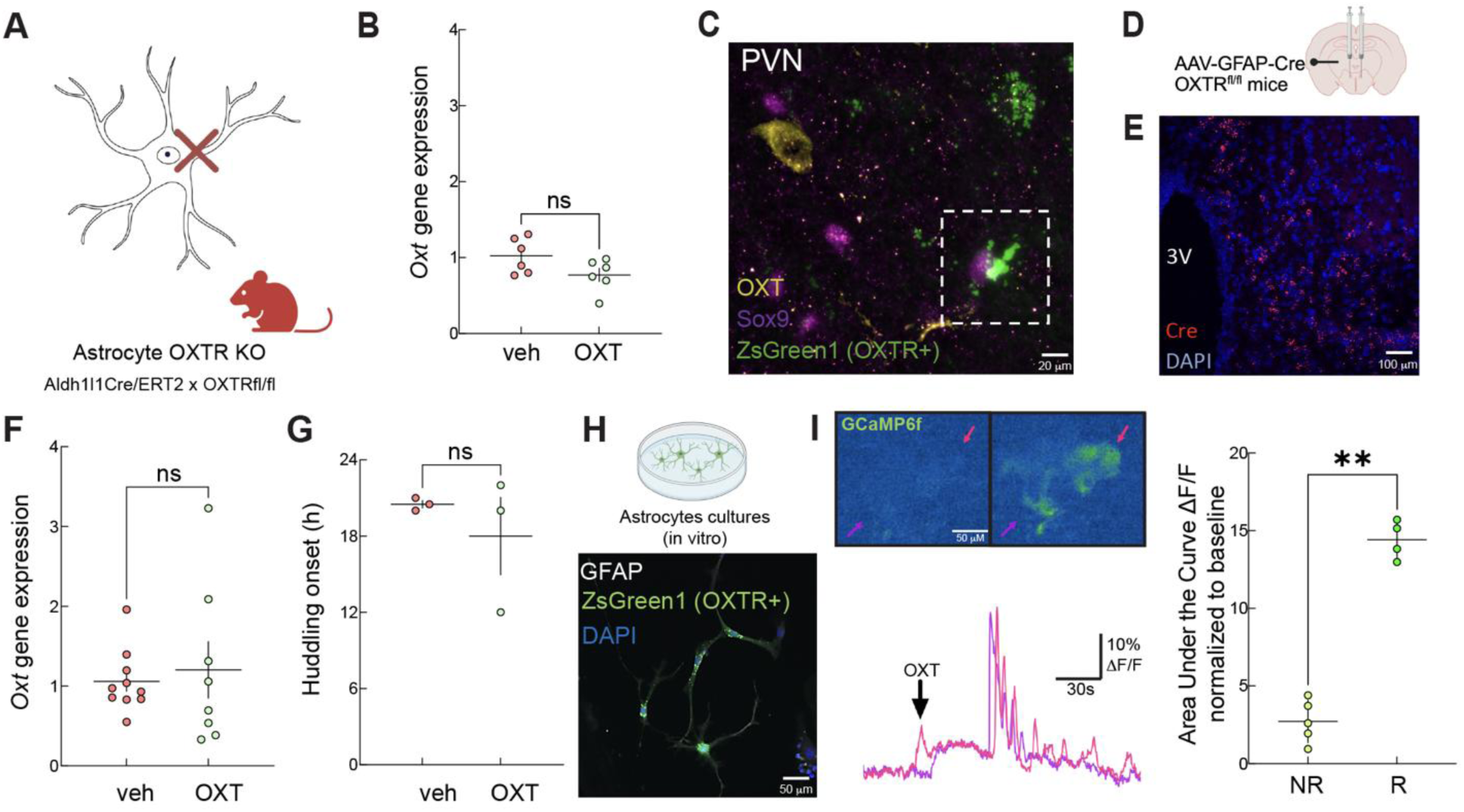
Cellular mediators of the oxytocin positive feedback loop. (**A**) OXTR knock-out strategy in astrocytes and (**B**) qPCR quantification of hypothalamic oxytocin mRNA levels. (**C**) Confirmation of OXTR-expressing (ZsGreen1) astrocytes (Sox9) surrounding oxytocin neurons (yellow) in the PVN of OXTR-Cre x Ai6 reporter mice. (**D**) Bilateral injection of an AAV-GFAP-Cre virus to locally knock-out OXTRs in PVN astrocytes and (**E**) confirmation of Cre (red) expression with RNAscope. (**F**) Quantification of hypothalamic oxytocin mRNA levels and (**G**) latency to start huddling upon reunion in PVN astrocytes OXTR knock-out mice. Each dot represents a mouse or a mouse pair in reunion experiments. Error bars represent SEM. (**H**) *In vitro* studies using OXTR-expressing (ZsGreen1) astrocyte (GFAP) cultures derived from OXTR-Cre x Ai6 reporter mice. (**I**) Example traces of oxytocin-induced calcium responses measured using an AAV-gfaABC1D-cyto-GCaMP6f virus and quantification of the area under the ΔF/F curve after oxytocin treatment normalized to baseline levels. NR: non-responders, R: responders. Each dot represents a cell. Error bars represent SD. Unpaired two-tailed Student’s t-test, **p<0.01, ns: not significant.

We next asked if this was locally mediated by PVN astrocytes. We confirmed OXTR expression in PVN astrocytes by crossing OXTR-Cre and Ai6 mice (**Fig. 3C**), then injected an AAV-GFAP-Cre virus bilaterally in the PVN of OXTR^fl/fl^ mice (**Fig. 3D,E**) to knock-out OXTRs only in PVN astrocytes. Mice lacking OXTRs in PVN astrocytes were no longer capable of upregulating oxytocin expression in response to exogenous oxytocin (**Fig. 3F**, p=0.6734). Moreover, isolated mice lacking OXTRs in PVN astrocytes were no longer rescued by intranasal oxytocin in terms of their behavioral phenotype, presenting huddling onset times comparable to untreated isolated mice (**Fig. 3G**, p=0.4610). These results show that astrocytes are the mediators of a local positive feedback loop for oxytocin expression in the PVN, and suggest that the behavioral effects of exogenous oxytocin are mediated by boosting the endogenous oxytocin system.

To understand molecular mediators of this positive feedback loop promoting oxytocin release, we next performed experiments in primary astrocyte cultures *in vitro*. We first confirmed OXTR expression in cultured astrocytes derived from OXTR-Cre x Ai6 reporter mice (**Fig. 3H**). To test the functionality of these receptors, we performed live calcium imaging in astrocyte cultures infected with an AAV-gfaABC1D-cyto-GCaMP6f virus. A subpopulation of astrocytes in culture expressed OXTRs and functionally responded to oxytocin (**Fig. 3I**, p<0.0001).

### Retinoic acid upregulates oxytocin expression

To characterize the cellular changes induced by oxytocin signaling in astrocytes at a molecular level, we exposed astrocyte cultures to 10 μM oxytocin added to the culture media 3 and 24 hours before processing the sample for transcriptomic analysis. We performed bulk RNA-seq of cultured astrocytes exposed to oxytocin and found a clear distinction between samples (**Fig. 4A**). Analysis of differentially expressed genes revealed Aldh1a1 as one of the most highly upregulated genes 24 hours after oxytocin treatment (**Fig. 4B**). The effect of oxytocin on Aldh1a1 is time-dependent and peaks at 24 hours (**Fig. 4C**).

**Fig. 4.**
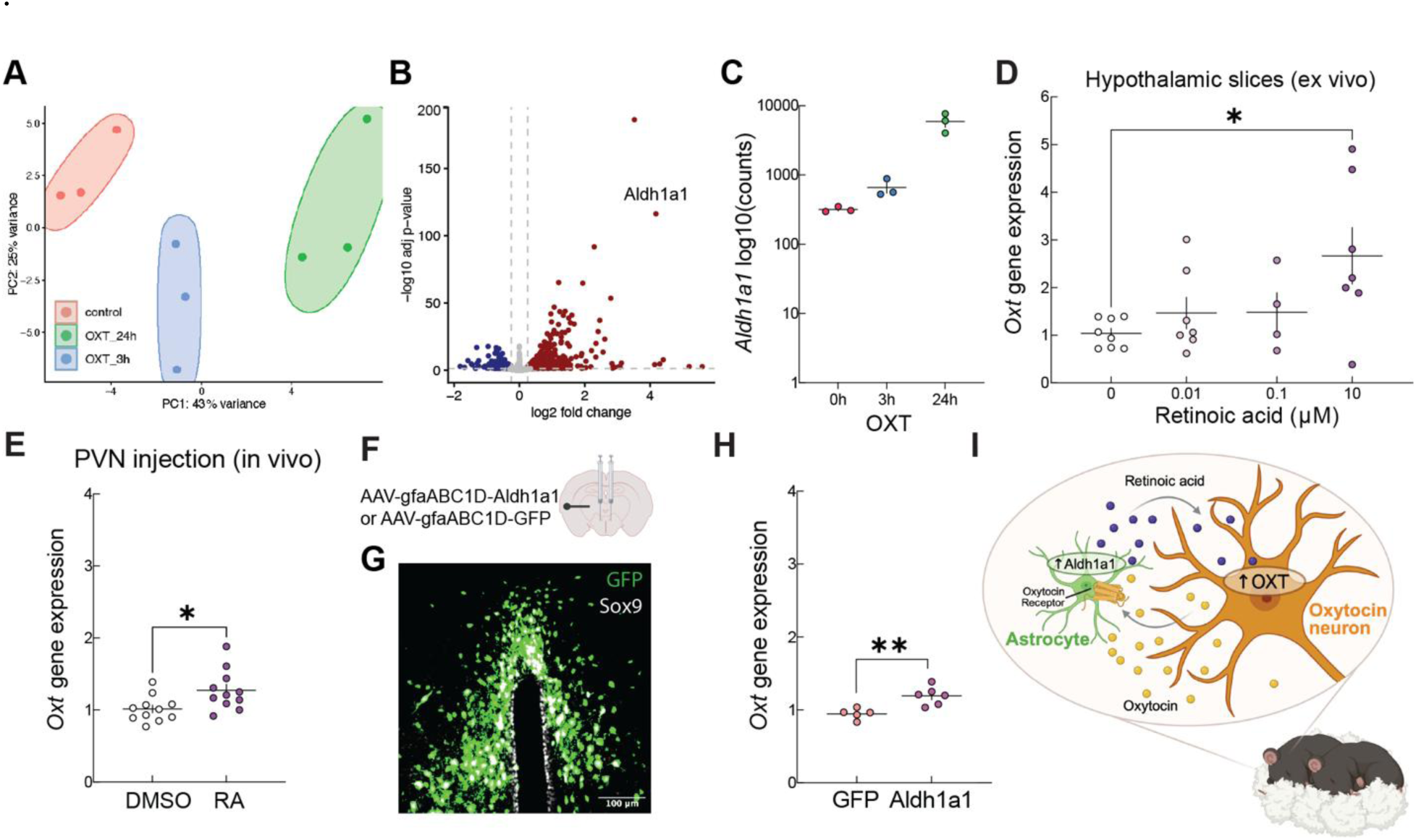
Molecular mediators of the oxytocin positive feedback loop. (**A** to **C**) Effects of oxytocin on astrocytes *in vitro*. (**A**) Principal component analysis (PCA) of controls and oxytocin (3 and 24 hours) treated astrocytes. (**B**) Differentially expressed genes (downregulated, blue and upregulated, red) in response to 24 hours of oxytocin treatment. (**C**) Time-dependent effects of oxytocin on Aldh1a1 (retinoic acid-synthesizing enzyme) expression. (**D**) Dose-dependent effects of 4 hours of retinoic acid exposure on oxytocin expression *ex vivo* in hypothalamic slices. One-way ANOVA. (**E**) *In vivo* effects after 6 hours of bilateral injections of 50 μM retinoic acid (or same dilution of DMSO as control) in the PVN on hypothalamic oxytocin mRNA levels. (**F**) Bilateral injection of an AAV-gfaABC1D-Aldh1a1 (AAV-gfaABC1D-GFP as control) virus to locally overexpress Aldh1a1 in PVN astrocytes and (**G**) confirmation of transgene (GFP) expression in astrocytes (Sox9). (**H**) *In vivo* effects of PVN astrocyte Aldh1a1 overexpression on hypothalamic oxytocin mRNA levels. (**I**) Schematic summarizing the cellular and molecular mediators of an oxytocin positive feedback loop that modulates social behaviors. Each dot represents an independent culture or a mouse. All error bars represent SEM. Unpaired two-tailed Student’s t-test, *p<0.05, **p<0.01.

Given that Aldh1a1 is an enzyme that catalyzes the last step of the retinoic acid biosynthesis pathway, we next studied whether retinoic acid could be increasing oxytocin expression. We tested the dose-dependent effects of retinoic acid *ex vivo* using hypothalamic slices. We found that retinoic acid induced a dose-dependent increase in oxytocin expression in this model after 4 hours of treatment (**Fig. 4D**, p=0.0326). Based on this data, we tested the effects of bilateral injections of 50 μM retinoic acid in the PVN *in vivo.* We found that after 6 hours, retinoic acid locally increases oxytocin expression compared to vehicle injected controls (**Fig. 4E**, p=0.0173).

Next, we tested whether Aldh1a1 expression upregulation in PVN astrocytes is sufficient to upregulate oxytocin expression. To do so, we injected an AAV-gfaABC1D-Aldh1a1 virus bilaterally in the PVN of wild-type mice, and AAV-gfaABC1D-GFP as control (**Fig. 4F,G**). We quantified oxytocin mRNA levels using qPCR and found that overexpression of Aldh1a1 in PVN astrocytes is sufficient to upregulate oxytocin expression (**Fig. 4H**, p=0.0041). In summary, this work shows that exogenous oxytocin delivery 1) increases endogenous oxytocin production through an astrocyte-dependent mechanism and 2) facilitates huddling upon reunion in socially isolated mice (**Fig. 4I**)

## Discussion

Social interactions have been proposed as a physiological need (*4, 17*). Social deprivation, induced by social isolation in mice, triggers opposite behavioral responses depending on the duration of the isolation period. While acute social isolation increases social seeking and prosocial behaviors, chronic deprivation leads to antisocial phenotypes (*17, 18*). Recent work from the Dulac laboratory has shown that the increase in social seeking induced after short periods of social isolation is mediated at least in part by oxytocin (*4*). In this study, we found that long-term social deprivation delays affiliative behaviors and downregulates oxytocin production in male mice, which can be prevented by exogenous oxytocin. While this effect of oxytocin-mediated upregulation of its own expression has been reported before in autism (*19–21*) and Alzheimer’s disease (*22*) mouse models, the cellular and molecular mediators of this effect have not been described before. We identified astrocytes as mediators of the loop, which appears to be necessary for the behavioral effects of exogenous oxytocin. This highlights the importance of assessing the functionality of this loop in oxytocin-based therapies.

While oxytocin signaling has mostly been studied in projecting areas, it can also be released locally through a somatodendritic mechanism (*23–25*). This allows surrounding cells to sense and respond to it. Hypothalamic astrocytes have previously been shown to transcriptionally respond to oxytocin to mediate anxiolytic behaviors (*26*) and a correlation between calcium responses in PVN oxytocin neurons and astrocytes has also been described during social interactions (*27*). This evidence, in line with our results, suggests that PVN astrocytes might be key players in the behavioral responses mediated by oxytocin. Moreover, a local effect of oxytocin on astrocytes has also been observed in the supraoptic nucleus (SON) - another hypothalamic nucleus containing oxytocin neurons.

During lactation, OXTR activation in astrocytes induces the retraction of their processes to facilitate oxytocin neuronal activity (*28, 29*). Moreover, oxytocin’s effect on astrocytes has also been described in projecting areas. Work from the Charlet laboratory shows that oxytocin signaling in amygdala astrocytes modulates neuronal activity and behavior (*12*). Characterizing the effects of oxytocin on neuronal and non-neuronal cell types in a brain-region specific manner is necessary to understand how this neuropeptide controls neuronal changes and behavioral outputs.

Since this work was done in adult male mice, whether oxytocin treatment can help alleviate molecular and behavioral changes induced by social deprivation during development or at later stages in life remains to be elucidated. Moreover, the behavioral outputs driven by oxytocin are known to be sexually dimorphic (*30, 31*) and social isolation has been described to have sex-specific effects in mice (*32, 33*), which has also been described in humans (*3, 34*) and prairie voles (*35–37*). Therefore, the effects of chronic social isolation on endogenous oxytocin production and the functionality of this loop across sexes and species need to be further studied.

At the molecular level, while the OXTR is known to be a G protein-coupled receptor (GPCR) (*38*), the exact mechanisms triggered by astrocytes in response to oxytocin remain to be fully elucidated. It is known that OXTR activation in astrocytes induces an increase in calcium levels (*12, 39*) and can activate Gq and Gi/o proteins (*14, 40*). However, as shown here and by Wahis and collaborators (*12*), only a subpopulation of astrocytes expresses OXTRs. It is likely that the fact that astrocytes are connected by gap junctions to form a network, allows for the intracellular signal activated by oxytocin to propagate across cells within this network (*12*). However, how astrocyte networks respond to oxytocin remains an open question.

This work also highlights the relevance of neuropeptide signaling in non-neuronal cells, which has increasingly gained attention. Recent studies have described a role of astrocytes in responding to and modulating neurotransmitter dynamics to ultimately influence behavioral outputs. Two recent studies describe the relevance of astrocytes for dopamine regulation. Lazaridis and collaborators (*41*), propose a loop in which striatal astrocytes sense and modulate dopamine release and Guttenplan and collaborators (*13*) show that astrocytes sense and modulate dopamine neuronal responses in a state-dependent manner. Another recent study by Sánchez-Ruiz and collaborators (*42*) shows that dorsal raphe astrocytes modulate serotonergic neurons to control social behaviors.

In line with the emerging role of astrocytes in sensing and responding to neuromodulators, we identified the cellular and molecular mediators of a crosstalk mechanism between oxytocin neurons and astrocytes in the context of social isolation. While these findings underscore the importance of astrocyte–neuron interactions in social behaviors, the precise dynamics and signaling pathways of the oxytocin–astrocyte loop remain to be further studied. This framework will contribute toward a comprehensive understanding of how diverse brain cell types coordinate complex social behaviors.

## Materials and Methods

### Animals

All experiments follow NYU Grossman School of Medicine Institutional Animal Care and Use Committee (IACUC) approved protocols and National Institutes of Health (NIH) guidelines. Mice were kept in an animal facility at ∼22°C and ∼45% humidity with a standard light/dark cycle. Mice were kept in cages containing multiple animals and only single housed for a 4 week-long social isolation period starting at the age of two months. Wild-type experiments involve the use of C57BL/6N mice (Charles River). Oxytocin-Ires-Cre (Strain #024234), Aldh1l1-Cre/ERT2 (Strain #031008), OXTR^fl/fl^ (Strain #008471), OXTR-Cre (Strain #031303) and Ai6 (Strain #007906) mice were obtained from The Jackson Laboratory and bred at our animal facility. Aldh1l1-Cre/ERT2 mice were induced by 5 consecutive days of gavage with a solution of 20 mg/mL tamoxifen dissolved in corn oil.

### Surgeries

Mice were anesthetized using 1-1.5% isoflurane and all surgical procedures were done using a stereotaxic instrument. For viral injections: mice received bilateral injections of one of the following viruses purchased from Vector Builder: AAV-GFAP-Cre, AAV-gfaABC1D-Aldh1a1, AAV-gfaABC1D-GFP or AAV-hSyn-DIO-hM3D in the PVN (coordinates in mm: −0.71 A-P, ±0.12 M-L, −4.7 D-V). For osmotic pump implantations: 28-day osmotic pumps were purchased from Alzet (MODEL 1004), were pre-loaded with CNO (or vehicle as control) and subcutaneously implanted close to the scapulae.

### Long-term behavioral monitoring

Mice were reunited in a long-term behavioral monitoring box (*15*) at ∼22°C with a standard light/dark cycle. The reunion sessions had a duration of 24 hours. During this whole time, videos were recorded and latency to the onset of huddling (for at least 30 minutes) was manually quantified.

### qPCR

RNA was extracted from homogenized samples using a RNA isolation and purification kit (Qiagen). Purity was assessed by the 260/280 nm absorbance ratio. cDNA was synthesized from one μg mRNA using a QuantiTect Rev. Transcription Kit (Qiagen). qPCR analysis was performed using the Power SYBR kit (Applied Biosystems) on a StepOnePlus™ Real-Time PCR System. Fold changes in gene expression were calculated using cycle threshold (Ct) values and normalized to the mean value of controls. The following primers were used for oxytocin: Forward: GAGGAGAACTACCTGCCTTCG; Reverse: TCCCAGAAAGTGGGCTCAG; and for the control actin-β: Forward: 5′TGTGACGTTGACATCCGTAAA-3′; Reverse: 5′GTACTTGCGCTCAGGAGGAG-3′.

### Whole brain imaging

Mice were anesthetized and perfused with phosphate-buffered saline (PBS), followed by 4% paraformaldehyde in PBS. Brains were collected, postfixed in 4% paraformaldehyde in PBS overnight, and processed following the iDISCO protocol (*43, 44*). Immunostaining was performed using a rabbit-anti-oxytocin primary antibody (Phoenix Pharmaceuticals G-051-01, 1:200) and an Alexa Fluor® 647 AffiniPure F(ab’)₂ Fragment donkey-anti-Rabbit IgG (H+L) (Jackson Immuno Research 711-606-152, 1:800) secondary antibody. Brains were imaged in a light sheet microscope (Zeiss Z1) with 5x and 20x lenses acquired using Zen Black software (Zeiss). Images were stitched using Zen Blue software (Zeiss) and Imaris software was used for reconstruction and volume quantification.

### Chemogenetics

Oxy-Cre mice were injected bilaterally in the PVN with an AAV-hSyn-DIO-hM3D(Gq) virus at the age of 1.5 months. Two weeks after viral injection, mice were subcutaneously implanted with an Alzet osmotic pump for continuous delivery of CNO (or vehicle) during the 4 week-long isolation period.

### Oxytocin delivery

Mice were gently restrained manually to receive 10 μL of a solution containing 8 ng of oxytocin or vehicle in both nostrils, as previously described by Selles and collaborators (*22*). This procedure was repeated 3 times a week for 4 weeks.

### Immunostaining

For immunohistochemistry, mice were anesthetized and perfused with phosphate-buffered saline (PBS), followed by 4% paraformaldehyde in PBS. Brains were collected, postfixed in 4% paraformaldehyde in PBS overnight, cryoprotected using increasing concentrations of sucrose, frozen in dry ice and stored at −80°C. Brains were sectioned (40 μm) in a cryostat (Leica Microsystems), blocked for 2 hours with 0.1% Triton X-100 and 5% Normal Donkey Serum (NDS) in PBS at room temperature, and incubated overnight at 4°C with anti-mouse-anti-Oxytocin (Millipore MAB5296, 1:250) and/or rabbit-anti-Sox9 (Abcam ab185966, 1:200) primary antibodies diluted in blocking buffer. On the following day, sections were washed and incubated for 2 hours at room temperature with Alexa Fluor 647-, 555-, or 488-conjugated secondary antibodies (Abcam, 1:500) and mounted with Prolong with DAPI (Invitrogen).

For immunocytochemistry astrocyte cultures were fixed using 4% paraformaldehyde in PBS 10 minutes, permeabilized with 0.1% Triton X-100 in PBS for 10 minutes, blocked using 10% NDS in PBS for 1h at room temperature and incubated overnight at 4°C with goat-anti-GFAP (Abcam ab53554, 1:500) in blocking buffer. Alexa-conjugated secondary donkey-anti-goat (Abcam) were prepared 1:1,000 in 1% NDS and incubated for 2 hours at room temperature. Cells were washed 3 times with PBS, mounted in Prolong with DAPI (Invitrogen) and imaged in a Zeiss LSM 800 confocal microscope. All images were acquired using a Zeiss LSM 800 confocal microscope with 20x or 63x objectives and processed using FIJI.

### RNAscope in situ hybridization

Brains were fresh frozen and processes following the RNAscope Multiplex Fluorescent Kit v2 Assay from Bio-Techne using a Cre probe to determine the presence of GFAP-driven expression of the enzyme in the PVN. All images were acquired using a Zeiss LSM 800 confocal microscope with a 20x objective and processed using FIJI (NIH).

### Astrocyte cultures

Mouse cortical astrocytes were purified from day 0-2 pups from wild-type or OXTR-Cre x Ai6 reporter mice, as previously described by Clayton and collaborators (35). Cells were cultured at 37°C and 10% CO₂. Once cells were around 70% confluent (5-7 days), cultures were dissociated and stored frozen. When used for experiments, cells were thawed and grown in basal medium supplemented with HBEGF (5 ng/mL) until they reach 70% confluence. Astrocyte cultures were processed for immunostaining or treated with 10 μM oxytocin or vehicle for imaging or sequencing experiments. For live calcium imaging oxytocin was directly added to the media during the imaging session and for bulk RNA-sequencing oxytocin was added to the media 3 or 24 hours before sample collection.

### Live calcium imaging

Astrocyte cultures were infected with an AAV-gfaABC1D-cyto-GCaMP6f virus 5 days before imaging. Fluorescence changes in response to oxytocin were imaged live on a spinning disk confocal (CrestOptics X-LIGHT V3 Confocal on Nikon Ti2) with controlled temperature and CO2 levels. Images were processed and quantified using FIJI (NIH).

### RNA sequencing

RNA was obtained from astrocyte cultures using a RNA isolation and purification kit (Qiagen). Isolated RNA was collected and analyzed for integrity with an Agilent Bioanalyzer. Libraries were prepared using an Automated stranded RNA-seq library prep with polyA selection and sequenced on a NovaSeqX 10B 100 cycle flow cells at the Genome Technology Center at NYU Grossman School of Medicine. FastQ files were run through fastqc and trimgalore. FastQ files were pulled and aligned using salmon and GENCODE reference mouse genome (gvM23 GRCm38.p6). Read counts were analyzed through the DESeq2 (1.44.0) and limma (3.60.6).

### *Ex vivo* hypothalamic slices

Hypothalamic tissue from 2-3 months old C57Bl/6 mice was dissected and sectioned in 400 μm slices using a McIlwain Tissue Chopper. Slices were incubated at 37°C and 5% CO₂ for 1 hour in artificial cerebrospinal fluid (aCSF) for recovery and treated with retinoic acid (0, 0.01, 0.1 or 10 μM) solubilized in DMSO and dissolved in aCSF. Samples were collected for qPCR analysis of oxytocin mRNA levels after 4 hours of treatment.

## Acknowledgments

We thank the Microscopy Core - particularly Michael Cammer and Joseph Sall, the Genotyping Core Laboratory, the Genome Technology Center, the Rodent Behavior Laboratory and the Small Instrument Fleet at New York University (NYU) Langone for experimental and technical support. We thank Theodore Fisher for guidance with RNAscope experiments, Luisa Schuster for her contribution to the behavioral experiments, the Lin lab at NYU for their contribution with reagents, and Jessica Minder, Catalina Varela, Jesus Morales Santos, Kirsten Tiangco and Sanjna Battepati.

## Author contributions

M.C.S. designed research; M.V.C. and R.C.F. supervised the work; M.C.S., M.L.C., F.L. and A.A. performed research; M.C.S. wrote the paper; M.C.S., M.L.C., F.L., A.A., S.A.L., R.C.F. and M.V.C. edited the paper.

## References

1. D. Wei, V. Talwar, D. Lin, Neural circuits of social behaviors: Innate yet flexible. Neuron 109, 1600–1620 (2021).

2. P. Chen, W. Hong, Neural Circuit Mechanisms of Social Behavior. Neuron 98, 16–30 (2018).

3. in Our Epidemic of Loneliness and Isolation: The U.S. Surgeon General’s Advisory on the Healing Effects of Social Connection and Community. (Washington (DC), 2023).

4. D. Liu et al., A hypothalamic circuit underlying the dynamic control of social homeostasis. Nature 640, 1000–1010 (2025).

5. R. C. Froemke, L. J. Young, Oxytocin, Neural Plasticity, and Social Behavior. Annu Rev Neurosci 44, 359–381 (2021).

6. Y. Tang et al., Social touch promotes interfemale communication via activation of parvocellular oxytocin neurons. Nat Neurosci 23, 1125–1137 (2020).

7. R. Triana-Del Rio et al., The modulation of emotional and social behaviors by oxytocin signaling in limbic network. Front Mol Neurosci 15, 1002846 (2022).

8. L. J. Young, Z. Wang, The neurobiology of pair bonding. Nat Neurosci 7, 1048–1054 (2004).

9. R. Menon, I. D. Neumann, Detection, processing and reinforcement of social cues: regulation by the oxytocin system. Nat Rev Neurosci 24, 761–777 (2023).

10. M. Knoop et al., The Role of Oxytocin in Abnormal Brain Development: Effect on Glial Cells and Neuroinflammation. Cells 11, (2022).

11. R. Patani, G. E. Hardingham, S. A. Liddelow, Functional roles of reactive astrocytes in neuroinflammation and neurodegeneration. Nat Rev Neurol 19, 395–409 (2023).

12. J. Wahis et al., Astrocytes mediate the effect of oxytocin in the central amygdala on neuronal activity and affective states in rodents. Nat Neurosci 24, 529–541 (2021).

13. K. A. Guttenplan et al., GPCR signaling gates astrocyte responsiveness to neurotransmitters and control of neuronal activity. Science 388, 763–768 (2025).

14. A. Baudon, E. Clauss Creusot, F. Althammer, C. P. Schaaf, A. Charlet, Emerging role of astrocytes in oxytocin-mediated control of neural circuits and brain functions. Prog Neurobiol 217, 102328 (2022).

15. L. Schuster et al. (bioRxiv, 2025).

16. K. A. Lyon, N. J. Allen, From Synapses to Circuits, Astrocytes Regulate Behavior. Front Neural Circuits 15, 786293 (2021).

17. N. Padilla-Coreano, K. M. Tye, M. Zelikowsky, Dynamic influences on the neural encoding of social valence. Nat Rev Neurosci 23, 535–550 (2022).

18. C. R. Lee, A. Chen, K. M. Tye, The neural circuitry of social homeostasis: Consequences of acute versus chronic social isolation. Cell 184, 2794–2795 (2021).

19. M. Tsurutani, T. Goto, M. Hagihara, S. Irie, K. Miyamichi, Selective vulnerability of parvocellular oxytocin neurons in social dysfunction. Nat Commun 15, 8661 (2024).

20. O. Penagarikano et al., Exogenous and evoked oxytocin restores social behavior in the Cntnap2 mouse model of autism. Sci Transl Med 7, 271ra278 (2015).

21. Y. C. Dai et al., Neonatal Oxytocin Treatment Ameliorates Autistic-Like Behaviors and Oxytocin Deficiency in Valproic Acid-Induced Rat Model of Autism. Front Cell Neurosci 12, 355 (2018).

22. M. C. Selles et al., Oxytocin attenuates microglial activation and restores social and non-social memory in APP/PS1 Alzheimer model mice. iScience 26, 106545 (2023).

23. M. Ludwig, G. Leng, Dendritic peptide release and peptide-dependent behaviours. Nat Rev Neurosci 7, 126–136 (2006).

24. F. Althammer, M. Eliava, V. Grinevich, Central and peripheral release of oxytocin: Relevance of neuroendocrine and neurotransmitter actions for physiology and behavior. Handb Clin Neurol 180, 25–44 (2021).

25. T. Qian et al., A genetically encoded sensor measures temporal oxytocin release from different neuronal compartments. Nat Biotechnol 41, 944–957 (2023).

26. C. P. Meinung et al., OXTR-mediated signaling in astrocytes contributes to anxiolysis. Mol Psychiatry 30, 2620–2634 (2025).

27. K. C. Sandoval, J. Rychlik, K. Y. Choe, Calcium Dynamics in Hypothalamic Paraventricular Oxytocin Neurons and Astrocytes Associated with Social and Stress Stimuli. eNeuro 12, (2025).

28. D. Li et al., Astrocytic Modulation of Supraoptic Oxytocin Neuronal Activity in Rat Dams with Pup-Deprivation at Different Stages of Lactation. Neurochem Res 46, 2601–2611 (2021).

29. Y. F. Wang, G. I. Hatton, Astrocytic plasticity and patterned oxytocin neuronal activity: dynamic interactions. J Neurosci 29, 1743–1754 (2009).

30. K. Li, M. Nakajima, I. Ibanez-Tallon, N. Heintz, A Cortical Circuit for Sexually Dimorphic Oxytocin-Dependent Anxiety Behaviors. Cell 167, 60–72 e11 (2016).

31. Y. Sofer et al., Sexually dimorphic oxytocin circuits drive intragroup social conflict and aggression in wild house mice. Nat Neurosci 27, 1565–1573 (2024).

32. L. Senst, D. Baimoukhametova, T. L. Sterley, J. S. Bains, Sexually dimorphic neuronal responses to social isolation. Elife 5, (2016).

33. M. Guo, C. F. Wu, W. Liu, J. Y. Yang, D. Chen, Sex difference in psychological behavior changes induced by long-term social isolation in mice. Prog Neuropsychopharmacol Biol Psychiatry 28, 115–121 (2004).

34. T. L. Procyshyn, J. Dupertuys, J. A. Bartz, Neuroimaging and behavioral evidence of sex-specific effects of oxytocin on human sociality. Trends Cogn Sci 28, 948–961 (2024).

35. K. Horie et al., Male, but not female, oxytocin receptor knockout prairie voles (Microtus ochrogaster) show impaired consolation behavior. Horm Behav 169, 105708 (2025).

36. K. L. Bales, C. S. Carter, Sex differences and developmental effects of oxytocin on aggression and social behavior in prairie voles (Microtus ochrogaster). Horm Behav 44, 178–184 (2003).

37. H. Pournajafi-Nazarloo et al., Effects of social isolation on mRNA expression for corticotrophin-releasing hormone receptors in prairie voles. Psychoneuroendocrinology 36, 780–789 (2011).

38. B. Jurek, I. D. Neumann, The Oxytocin Receptor: From Intracellular Signaling to Behavior. Physiol Rev 98, 1805–1908 (2018).

39. D. Di Scala-Guenot, D. Mouginot, M. T. Strosser, Increase of intracellular calcium induced by oxytocin in hypothalamic cultured astrocytes. Glia 11, 269–276 (1994).

40. A. Baudon et al., Stress induces oxytocin-Galphai-dependent remodeling of astrocytes to shape neuronal response in the amygdala. Nat Commun, (2025).

41. I. Lazaridis, G. Ahn, K. Hirokane, W. Choi, A. M. Graybiel. (bioRxiv, 2024).

42. A. Sánchez-Ruiz, C. González-Arias, L. Arancibia, S. Mederos, G. Perea. (Research Square, 2025).

43. N. Renier et al., iDISCO: a simple, rapid method to immunolabel large tissue samples for volume imaging. Cell 159, 896–910 (2014).

44. M. L. Cooper et al., Astrocytes in the mouse brain respond bilaterally to unilateral retinal neurodegeneration. Proc Natl Acad Sci U S A 122, e2418249122 (2025).

